# Single neuron analysis of aging associated changes in learning reveals progressive impairments in transcriptional plasticity

**DOI:** 10.1101/2023.06.23.546336

**Authors:** Kerriann K Badal, Abhishek Sadhu, Carrie McCracken, Bindu L Raveendra, Sebastian Lozano-Villada, Amol C Shetty, Phillip Gillette, Yibo Zhao, Dustin Stommes, Lynne A Fieber, Michael C Schmale, Anup Mahurkar, Robert D Hawkins, Sathyanarayanan V Puthanveettil

## Abstract

Molecular mechanisms underlying aging associated impairments in learning and long-term memory storage are poorly understood. Here we leveraged the single identified motor neuron L7 in *Aplysia,* which mediates a form of non-associative learning, sensitization of the siphon-withdraw reflex, to assess the transcriptomic correlates of aging associated changes in learning. RNAseq analysis of the single L7 motor neuron isolated following short-term or long-term sensitization training of 8,10 and 12 months old *Aplysia,* corresponding to mature, late mature and senescent stages, has revealed progressive impairments in transcriptional plasticity during aging. Specifically, we observed modulation of the expression of multiple lncRNAs and mRNAs encoding transcription factors, regulators of translation, RNA methylation, and cytoskeletal rearrangements during learning and their deficits during aging. Our comparative gene expression analysis also revealed the recruitment of specific transcriptional changes in two other neurons, the motor neuron L11 and the giant cholinergic neuron R2 whose roles in long-term sensitization were previously not known. Taken together, our analyses establish cell type specific progressive impairments in the expression of learning- and memory-related components of the transcriptome during aging.

## INTRODUCTION

Advances in medicine have progressively extended human longevity but have amplified emerging health challenges, particularly aging-associated cognitive decline and developing debilitating diseases. Therefore challenges to modern medicine include understanding why and how we age and the possibility of reversing aging, which could significantly improve our lives. Decades of research using multiple animal models such as C. elegans (*1, 2*), Drosophila (*3, 4*), Aplysia (*5–7*) rodents (*8, 9*) and humans (*10, 11*) have identified several molecular and cellular changes underlying nervous system aging. These include changes in transcription, translation, the epigenome, and synaptic function and plasticity (*12–16*). Among these molecular changes, transcriptional changes are of particular interest because they set the stage for subsequent modifications of cellular signaling and intercellular communication. Furthermore, learning and long-term memory storage (LTM) require activation of gene expression changes (*17*) in specific neurons. This change in transcriptional state in response to learning, also described as “transcriptional plasticity”, or the ability of genes to change their expression when the environment changes (*18, 19*) is necessary for establishing learning and LTM.

Though aging-associated large-scale transcriptional and epigenetic changes in the nervous system have been described, we know very little about aging associated changes in individual neurons in a neural circuit during learning (*8, 20–22*). Currently available gene expression datasets on neuronal aging lack circuit-specific nor neuron-specific changes relevant for learning and memory storage. To address this knowledge gap, we exploited the advantages of identified neurons mediating learning in the sea slug *Aplysia Californica*, a neurobiological model for the cellular and molecular mechanisms of learning and long-term memory storage (LTM) (*17, 23–31*). Importantly, behavioral learning of the siphon withdrawal reflex (SWR) in *Aplysia* is well characterized (*65–67*). The SWR is a defensive reflex that undergoes both non-associative learning including sensitization and associative learning including conditioning (*32–36*). During sensitization responses (siphon withdrawals) elicited by weak test stimuli (a gentle tap to the siphon using a paint brush bristle) are augmented by training with a strong stimulus (electric shocks to the tail). A single electrical shock to the tail of Aplysia produces short-term sensitization or STS lasting several minutes whereas four spaced shocks to the tail produce long-term sensitization or LTS lasting several days.

To gain molecular insights into modulation of learning-relevant transcriptional changes during aging, we focused on sensitization, a form of non-associative learning, and assessed aging-associated changes in the expression of mRNAs and long-noncoding RNAs (lncRNAs) in L7 motor neuron (L7MN), a critical component of SWR circuitry. Briefly, we used RNAseq analyses to examine STS and LTS induced changes in the transcriptional landscape of L7 neurons at three ages. We first identified components of transcriptional plasticity of L7MN in 8 month old *Aplysia* and then compared that with L7MN isolated from 10 and 12 months old animals. Consistent with the previous literature on aging-associated behavioral changes in Aplysia (*37–40*), we identified deficiencies in LTS during aging. Our gene expression analyses revealed progressive impairments in both the coding and long-noncoding transcriptome of L7MN. We also identified shared and neuron specific learning and aging associated changes in gene expression in two other neurons (motor neuron L11 and Cholinergic neuron R2) whose roles in STS or LTS were not known.

## RESULTS

### Behavioral training showed impairments in long-term sensitization of siphon withdrawal reflex

Since LTM formation requires gene expression and new protein synthesis (*17, 23, 41-44*), we searched for transcriptomic alterations that are associated with aging related changes in LTS. To gain insights into the transcriptomic bases of aging associated impairments in LTS we sought to carry out total RNAseq analysis of L7 motor neurons (L7MN) from trained (STS or LTS) animals and untrained age matched controls across the age groups. The monosynaptic connection between the siphon sensory neurons and motor neuron L7 participates in learning and LTM of the siphon withdrawal reflex (*34, 36, 45*). Importantly, there is only one L7MN in the entire animal and it can be easily identified by its specific size and location in the abdominal ganglion.

We set up two cohorts of animals and from each cohort we measured gene expression and behavior changes in *Aplysia* 8 months (Age group1, sexually mature adults), 10 months (Age group 2, late mature) or 12 months old (Age group 3, senescent). *Aplysia* can live up to 12-14 months old in captivity under normal conditions of maintenance at the National Aplysia Resource Facility, although their life span can be prolonged by changing diet and temperature (*46*). Since sensitization of the siphon withdrawal reflex is a robust form of behavioral learning, from each cohort of animals we were able to use two animals per condition per age group for behavior assessments of long-term sensitization (LTS) with untrained animals as controls, and 8 animals per condition for single neuron isolations, which were carried out one hour after behavioral training. Measurements of the duration of siphon withdrawal 24 hours after LTS training compared to age matched controls showed that the duration of siphon withdrawal was significantly higher in 12 month old (Age group 3) animals suggesting impairments LTS during aging (Fig. 1, Supplementary Table S1A).

**Figure 1.**
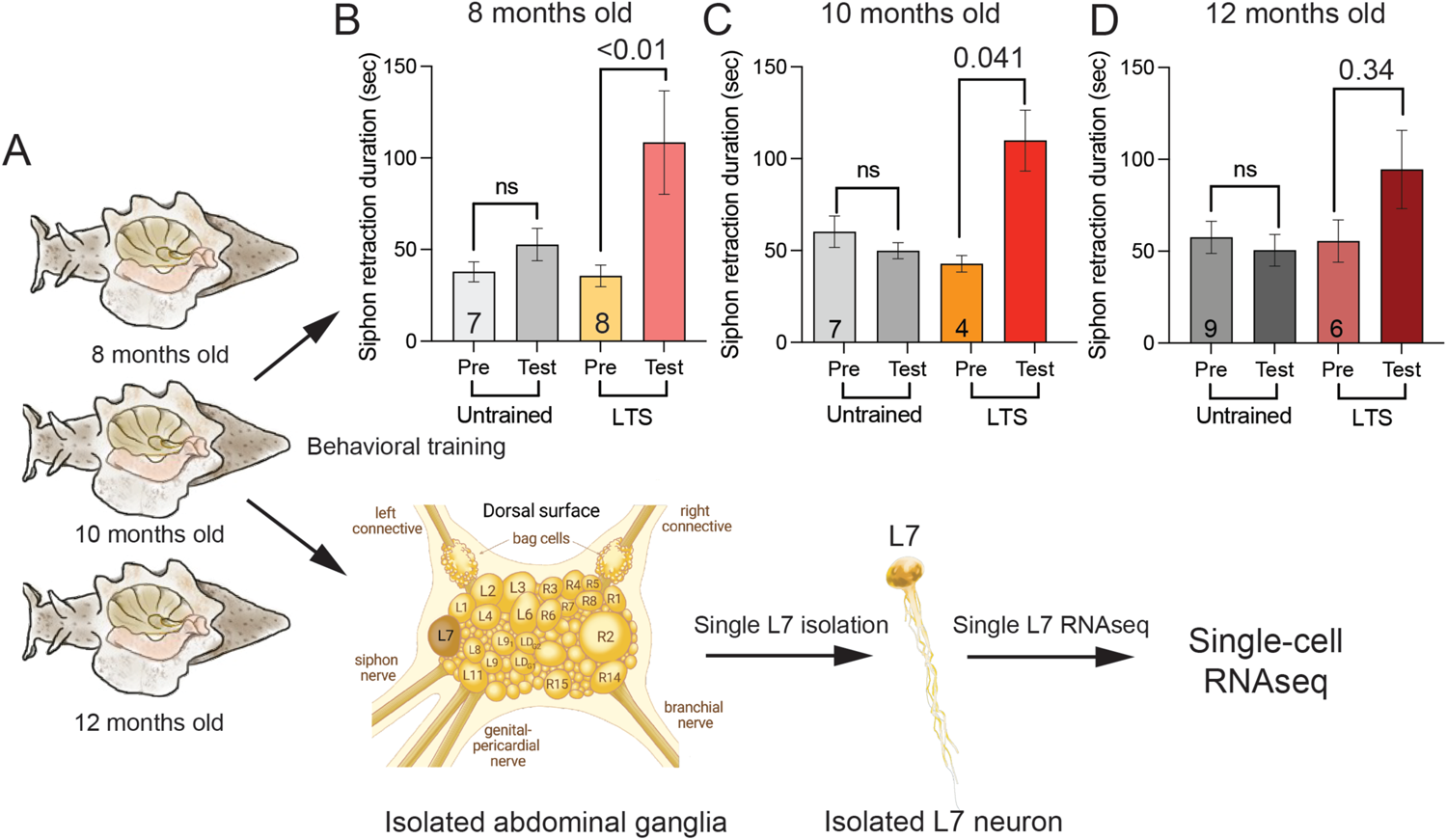
Overview of single neuron analysis of aging associated changes in learning. **A.** Schematic representation of the workflow for single L7 motor neuron (L7MN) isolation to RNAseq from trained (short-term and long-term sensitization) and untrained animals from 3 age groups. Group 1: 8 months, Group 2: 10 months, and Group 3: 12 months old., **B–D.** Bar graphs showing the average duration or latency of siphon withdrawal from the stimulus to the time the siphon begins to relax before (Pre) and after (Test, 24 hour after) long-term sensitization (LTS) training in three age groups. The number of animals used for analysis is shown in the bar graphs. NS: non-significant. One Way ANOVA, Error bars are SEM.

### Short-term and long-term sensitization training induces specific changes in long-noncoding and coding transcriptomes

Because STS and LTS training of SWR produces short-term and long-term memory of tail shock, to assess how aging impacts these two temporally distinct forms of memories, we first carried out STS and LTS training of 8 months old animals and isolated L7MN for total RNAseq analysis. L7MN RNAs from age matched untrained animals were used for comparisons This analysis identified 1314 unique RNA sequences including 82 lncRNAs in from L7MNs isolated from STS- and LTS-trained animals (Supplementary Table S1B). Differential expression analysis suggested 629, 364, and 706 differentially expressed genes (DEGs) between STS vs control, LTS vs control and LTS vs STS respectively covering 2.15%, 1.24% and 2.41% of all annotated genes (AplCal3.0; GCF_000002075.1; total 29270 transcripts) (Fig 2. A–J). Venn diagrams shown in Figure 2 suggest that STS and LTS training paradigms alter transcription of specific populations of mRNAs and lncRNAs.

**Figure 2.**
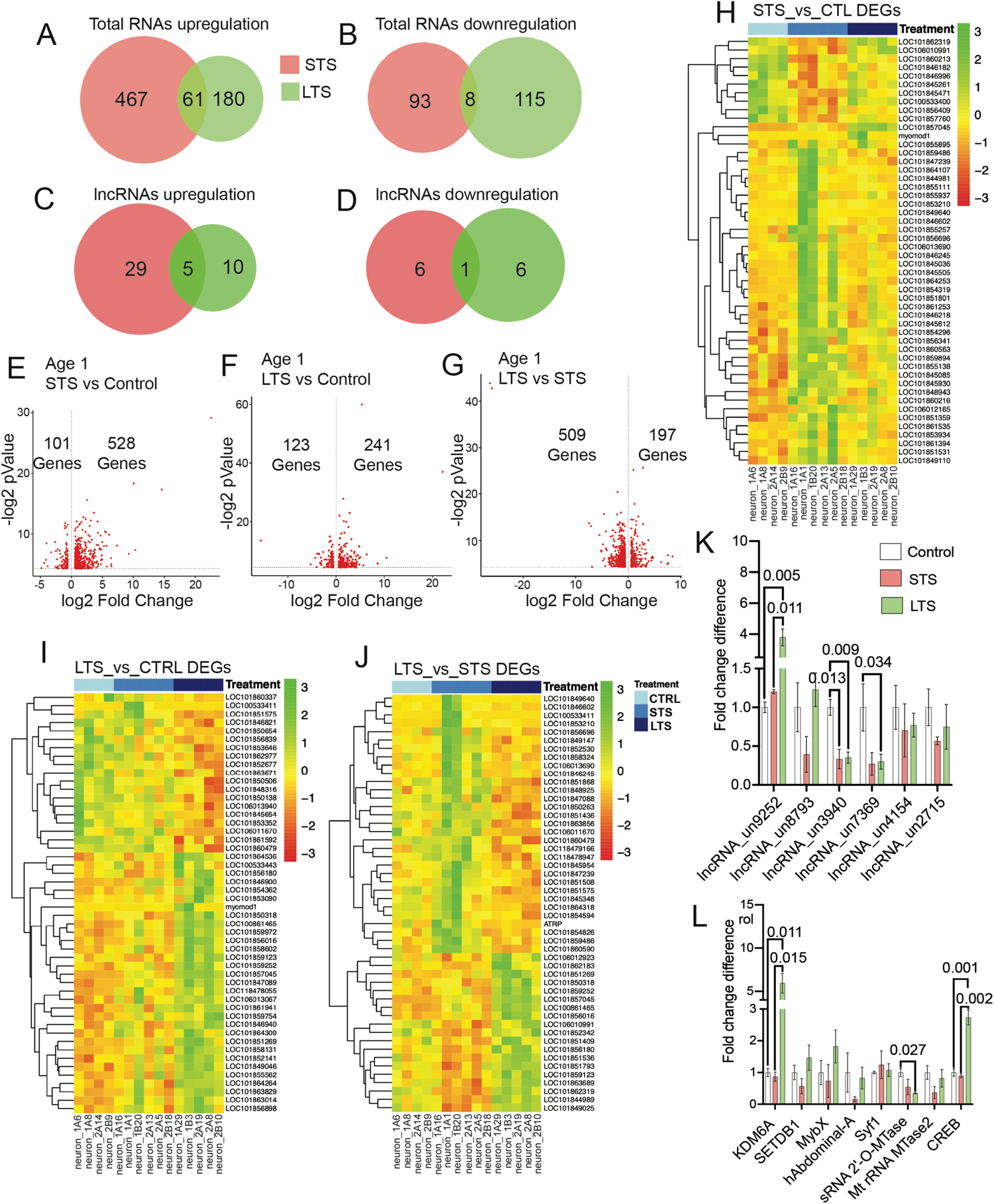
RNAseq analysis of L7MN reveals specific changes in the expression of mRNAs and long-noncoding RNAs (lncRNAs) following STS and LTS training. Venn diagrams showing (the numbers indicate unique and common DEGs) differentially expressed genes (DEGs) in 8 months old animals. **A.** upregulated **B.** downregulated in response to short-term sensitization, STS and long-term sensitization, LTS (*p*Value <0.05; The numbers indicate unique and common DEGs). Venn diagrams showing differentially expressed lncRNAs **C.** upregulated **D.** downregulated in response to STS and LTS (*p*Value <0.05). Differentially expressed genes are ranked in a volcano plot according to their statistical -log2 p-value (y-axis) and their relative abundance ratio (log2 fold change) between up- and downregulated (x-axis). Red dots indicate significantly regulated genes (false discovery rate, <0.01; s0 = 1; *p*Value <0.05). Volcano plots of **E.** Control versus STS DEGs, **F.** Control versus LTS DEGs, **G.** LTS versus STS DEGs. Heatmaps showing the normalized and scaled expression values of the top 50 differentially expressed genes when ranked by p-value. The color gradient from green to red represents high to low expression levels across the samples. The genes are ordered by hierarchical clustering using euclidean distance and complete clustering method while the samples are ordered by condition, **H.** STS versus Control, **I.** LTS versus Control, **J.** LTS versus STS. qPCR validation of selected candidates from RNAseq data, **K.** lncRNAs, **L.** mRNAs. Relative gene expression levels are exhibited as the mean fold change, with error bars showing the SEM. One-way ANOVA followed by Tukey’s *post hoc* test. N=4, p values are shown in the bar graphs.

Importantly these DEGs are involved in strengthening of the existing synaptic connections. For example, STS training has led to upregulation of RNA transcripts, among them ∼6.2% are lncRNAs, among mRNAs, ∼5% related to synapse function, ∼3.2% are related to transcription/translation, ∼6% are kinases or phosphatases. Examples of STS specific regulated mRNAs include multi drug resistance-associated protein 1 (LOC101849640), eukaryotic translation initiation factor 5A (LOC101854486) and other translation initiation factors (LOC101854486, LOC101853210, LOC101850385), adenylate cyclase (LOC101859667, LOC101860364), adhesion G protein-coupled receptor L1 (LOC101862738), several cytochrome P450s (LOC101853171, LOC101851093, LOC101852221, LOC101861801, LOC101864543), DNA polymerase alpha catalytic subunit isoform (LOC101861408), FMRFamide receptor (LOC101852874, LOC101859894), and syntaxin-7 (LOC101861711). Similarly we identified upregulation of 34 lncRNAs of which 5 are commonly upregulated in LTS and STS (Supplementary table S1B). By contrast LTS training resulted in the upregulation of 241 transcripts relative to long-term memory formation/ consolidation, ∼5.5% are lncRNA, ∼10.5% are synaptic proteins, ∼8.8% are involved in transcription or translation and ∼4.4% are kinase or phosphatases. cAMP-responsive element-binding protein (CREB, LOC100861465), CREB3 regulatory factor isoform (LOC101858375), probable G-protein coupled receptor 83 (LOC101855582), calcium/calmodulin-dependent protein kinase type 1B (LOC101845492), G-protein coupled receptor GRL101 (LOC101858538), glutamate receptor 2 (LOC100533395), calcium/calmodulin-dependent protein kinase type 1B (LOC101845492), acetylcholine receptor subunit alpha-type acr-16-like precursor (LOC106012547), lysine-specific demethylase 6A (LOC101851269) were found upregulated upon LTS, along with a total of 15 lncRNAs (Supplementary Table S1B). Moreover, when comparing LTS to STS, we found long-term memory related DEGs such as sonic hedgehog protein A (LOC101856180), which has been shown to be activated in the rodent Amygdala during learning (*47*), vacuolar protein sorting-associated protein isoform (LOC101859486), which encodes for proteins related to MIT: domain contained within Microtubule Interacting and Trafficking molecules (NIH Gene database) and cAMP responsive element-binding protein (CREB1) (LOC100861465).

We identified several commonly upregulated DEGs during STS and LTS (example: muscle contracting myomod1 (myomodulin neuropeptides 1 precursor), a response associated with escape behavior due to sensitization(*48*) as well as differentially modulated genes. For example. different isoforms of FMRFamide receptor, mucin-5AC, pedal peptide 2, snRNA_U4 spliceosomal RNA, upregulated during STS, were found to be downregulated during LTS. Similarly, isoforms of calcyphosin-like protein, multiple epidermal growth factor-like domains protein 10 downregulated upon STS were found upregulated in LTS. Several lncRNAs were also found to downregulated in STS and LTS. Examples of downregulated transcripts in LTS include dual specificity protein phosphatase 14 (LOC101848709), syntenin-1 (LOC101854063), small RNA 2’’-O-methyltransferase (LOC101850673). Taken together, these results show that STS and LTS alter expression of specific sets of mRNAs and lncRNAs in L7MN. Several known genes involved in memory processes (examples: CREB, CaMK II, lysine demethylase) are upregulated in L7MN following LTS unlike STS. While the role of lncRNAs in these STS and LTS are not known, our analysis suggest that lncRNAs are targets of transcriptional modulation relevant for learning and LTM in *Aplysia*.

### Validation of differential regulation of lncRNAs and mRNAs by STS and LTS

We next validated our RNAseq data in L7MN isolated following another round of STS and LTS training. Based on the fold enrichments and known functions in learning and LTM relevant process, we selected eight mRNAs (five upregulated and three downregulated) and analyzed the gene expression levels by qPCR and six lncRNAs (two upregulated, and four downregulated). All the lncRNAs we examined were not examined previously. Aplysia 18S rRNA gene was used to normalize the gene expression levels. A list of primers used in the study has been provided in the Supplementary Table S1B.

Consistent with the RNAseq data, we found that following LTS lncRNA_un9252 showed ∼3.8-fold (*p*<0.01) increase in gene expression level compared to control and ∼3-fold (*p*<0.05) higher compared to STS (Fig 2K); lncRNA_un3940 and lncRNA_un7369 showed ∼2.8-fold and ∼6-fold decrease compared to control. Figure 2L summarizes the gene expression levels of the selected mRNA candidates. Analyses showed that following LTS training, KDM6A showed ∼6-fold upregulation (*p*<0.05) in expression level compared to control and sRNA 2’-O-MTase level was ∼2.8-fold down regulation (*p*<0.05) compared to control. We observed enhancements in CREB level ∼2.7-fold (*p*<0.01), a previously characterized transcriptional activator essential for learning and LTM (*49*). These successful validation by independent experiments further support that STS and LTS training produces specific changes in lncRNA and mRNA expressions in L7MN.

### DEG analysis of L7MN from 10 and 12 months old animals suggest progressive impairments in transcriptional plasticity

To understand the transcriptomic bases of aging-associated decline in LTS, we next analyzed the L7MN transcriptome from ten months (Age 2) and twelve-month-old (Age 3), trained animals. We identified 1317 and 1460 transcripts, including 52 and 75 lncRNA from the age group 2 and 3 respectively. Differential expression analysis of age group 2 suggested 747, 399, and 545 genes (Fig 3. A–F Supplementary Table S1C) were differentially expressed between STS vs control, LTS vs control and LTS vs STS respectively covering ∼2.54%, ∼1.36% and ∼1.86% of the annotated genes (AplCal3.0; GCF_000002075.1; total 29270 transcripts), and in age group 3 we observed 421, 1031 and 382 genes differentially expressed between STS vs control, LTS vs control and LTS vs STS respectively covering ∼1.43 %, ∼3.52% and ∼1.3% of the *Aplysia* genome (Fig 3. G–L; Supplementary S1C AplCal3.0; GCF_000002075.1; total 29270 transcripts).

**Figure 3.**
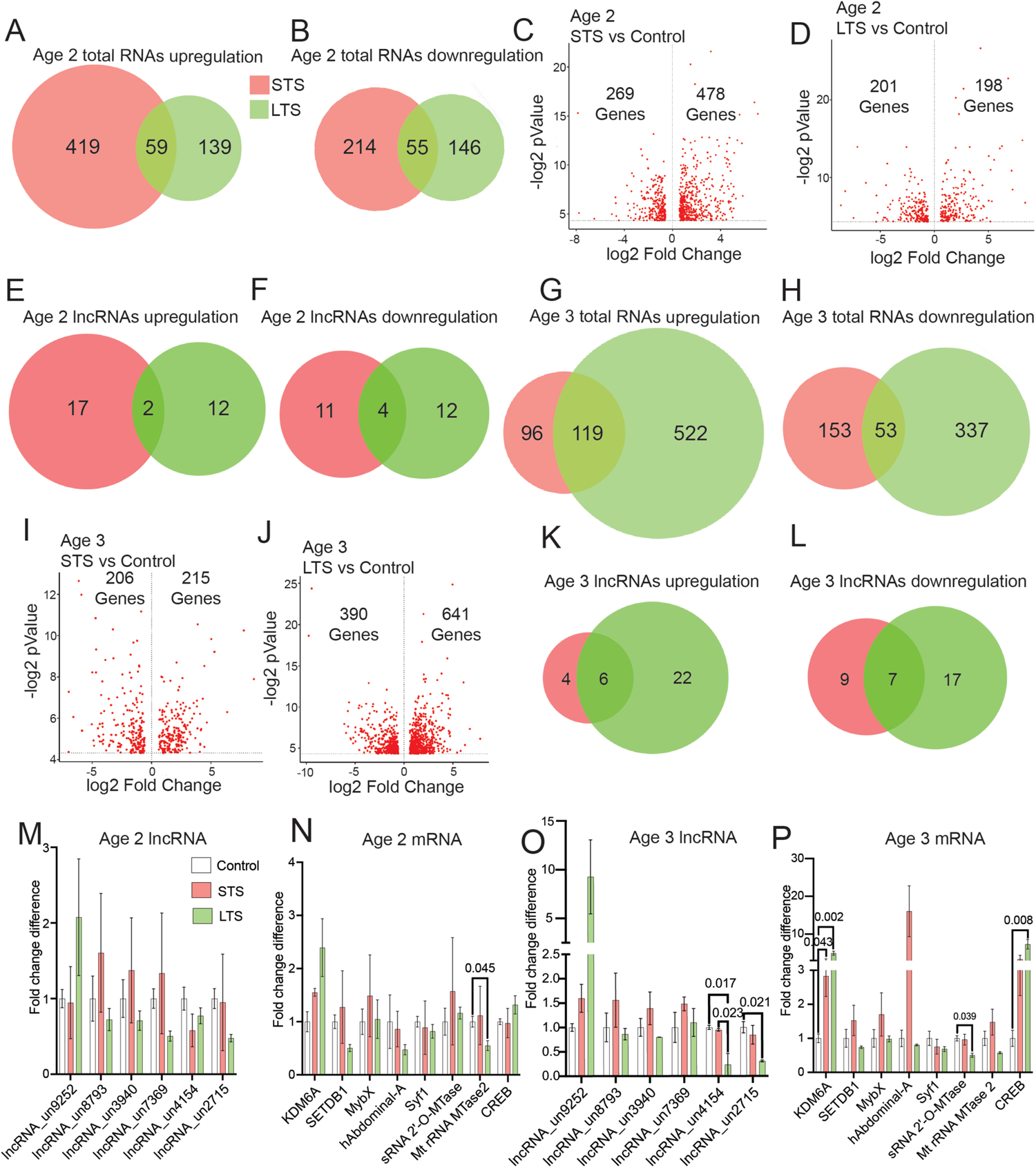
RNAseq analysis of L7MN from 10 and 12 months old Aplysia following STS and LTS training. Venn diagram showing Age 2 DEGs **A.** upregulated **B.** downregulated in response to STS and LTS (*p*Value <0.05). Differentially expressed genes are ranked in a volcano plot according to their statistical -log2 p-value (y-axis) and their relative abundance ratio (log2 fold change) between up- and downregulated (x-axis). Red dots indicate significantly regulated genes (false discovery rate, <0.01; s0 = 1; *p*Value <0.05). Volcano plots of **C.** Age 2 Control versus STS DEGs, **D.** Age 2 Control versus LTS DEGs. Venn diagram showing DEG lncRNAs **E.** upregulated **F.** downregulated in response to STS and LTS (*p*Value <0.05). Venn diagram showing Age 3 DEGs. **G.** upregulated **H.** downregulated in response to STS and LTS (Age 3; *p*Value <0.05). Volcano plots of **I.** Age 2 Control versus STS DEGs, **J.** Age 2 Control versus LTS DEGs. Venn diagram showing DEG lncRNAs **K.** upregulated **L.** downregulated in response to STS and LTS (Age 3; *p*Value <0.05). qPCR validation of the RNAseq data **M.** Age 2 lncRNAs, **N.** Age 2 mRNAs, **O.** Age 3 lncRNAs, **P.** Age 3 mRNAs. Relative gene expression levels are shown as the mean fold change, with error bars showing the SEM. One-way ANOVA followed by Tukey’s *post hoc* test. N=4, p values are shown in the bar graphs.

In Age 2 group during LTS, among the upregulated DEGs with synapse signaling were brain-specific homeobox/POU domain protein 3-like isoform (LOC101859216), calcium/calmodulin-dependent 3’,5’-cyclic nucleotide phosphodiesterase 1C (LOC106014171), GTP-binding protein RAD (LOC106014213), synapse-associated protein 1 (LOC101862979), potassium voltage-gated channel subfamily H member 1 (LOC101860035), FMRFamide receptor (LOC101850131); transcription/translation factors such as myb-like protein A (LOC101847950), transcription factor MYB120 (LOC101858259), zinc finger A20 and AN1 domain-containing stress-associated protein 9 (LOC106013912), forkhead box protein biniou (LOC101856898), Krueppel-like factor 8 (LOC101850318). In Age 3 group during LTS, we observed synaptic proteins such as cyclic AMP-dependent transcription factor ATF-7 (LOC101856191), serine/threonine-protein kinase fray2 (LOC106012574), cAMP responsive element-binding protein (LOC100861465), pannexin 2 (LOC100533356), ankyrin repeat domain-containing protein 17 (LOC101846593), metabotropic glutamate receptor 3 (LOC101863041), serine/threonine-protein kinases D1, fray2, pakF, RIO1, SIK3 (LOC101864159, LOC106012574, LOC101854275, LOC101845822, LOC101849046) etc.; transcription factors such as forkheadbox protein K2 and O (LOC101847706, LOC101847009), zinc finger protein 16, 271, 493, 628, 704, 708 (LOC101848424, LOC101851507, LOC101852427, LOC118477209, LOC101845787, LOC101860932, LOC101857232), transcription factor 20, MafF, Sox-10, Sp3, TFIIIB (LOC101853672, LOC101851728, LOC101847270, LOC101863331, LOC101855811). Interestingly we identified several epigenetic regulation related genes such as histone acetyltransferase KAT2A isoform (LOC101856257), histone-lysine N-methyltransferase ASH1L isoform (LOC101849998), GADD45 (LOC101853028, LOC101846541), KAT8 regulatory NSL complex subunit 1 (LOC101845902), N-acetyltransferase ESCO2 (LOC101857418), uncharacterized methyltransferase C25B8.09 (LOC101861999), beta-1,4-N-acetylgalactosaminyl transferase bre-4 (LOC101851752), threonylcarbamoyl-adenosine tRNA methylthio-transferase (LOC101863090).

Among the downregulated DEGs in Age 2 group, FMRFamide-related neuropeptides-like (LOC101851187), voltage-gated potassium channel subunit beta-2 (LOC101861756), synaptotagmin-1 (LOC101856907) and in Age 3 group, calmodulin (LOC101850552), voltage-dependent calcium channel subunit alpha-2/delta-3 (LOC101847330, LOC101847576), synapsin isoform 2.1 (LOC100533225), neuronal acetylcholine receptor subunit alpha/beta-4 (LOC101861149, LOC101845835) are some of the important genes involved in synapse function. Taken together, RNAseq analysis across the three age groups indicate escalating alterations in gene expression in L7MN.

### Validation of differential expression of candidate lncRNAs and mRNAs in 10 and 12 months old animals

Focusing on the same lncRNA and mRNA candidates that we studied in age group 1(8 months old), we next validated RNAseq data by qPCR analysis of L7MN isolated from ten-(Age 2) and twelve-months (Age 3) old animals. Fig 3. M shows no significant changes in the lncRNA levels were observed in Age 2 group. Upon LTS we found Mt rRNA MTase2 showed ∼3-fold (*p*<0.05) decrease in the expression level compared to untrained L7 (Fig 3. N).

In Age 3 animals, in accordance with the RNAseq data we observed that upon LTS lncRNA_un4154 and lncRNA_un2715 showed ∼4.3-fold (*p*<0.05) and ∼3.2-fold (*p*<0.05) decline in the expression level compared to control animals (Fig 3. O). Among analyzed mRNAs, in Age 2 group, KDM6A expression level was observed ∼2.8-fold (*p*<0.05) and ∼5-fold (*p*<0.01) increased upon STS and LTS than the control expression level (Fig 3. P). We also found that following LTS, the expression of sRNA 2’’-O-Mtase expression level declined ∼2-fold (*p*<0.05). CREB levels in Age 3, L7s was observed ∼6.4-fold (*p*<0.01) higher than the control level. Taken together these results suggest progressive decline in the expression of learning relevant lncRNAs and RNAs during aging. Furthermore, that multiple genes modulating nuclear function such as transcription and synapse function such as synaptic transmission are impacted in L7MN during aging.

### RNAseq data analysis showed changes in basal level gene expression across the age groups

To assess how aging impacts basal level transcriptome in L7MN, we compared RNAseq data of untrained animals across the three age groups. Fig 4. A–D shows the comparison of total RNA and lncRNAs up- and down-regulated DEGs in Venn diagrams. We found that 1326 transcripts were upregulated and 675 transcripts were downregulated in Age 2 vs Age 1 controls; 1188 transcripts upregulated, and 673 transcripts downregulated in Age 3 vs Age 1 controls; and 187 transcripts upregulated, and 227 transcripts were noted downregulated in Age 3 vs age 2 controls (Supplementary Table S1D). Fig 4. E–G represents the DEGs distribution in volcano plots. Taken together these results suggest that aging progressively impairs learning relevant transcriptome on L7MN.

**Figure 4.**
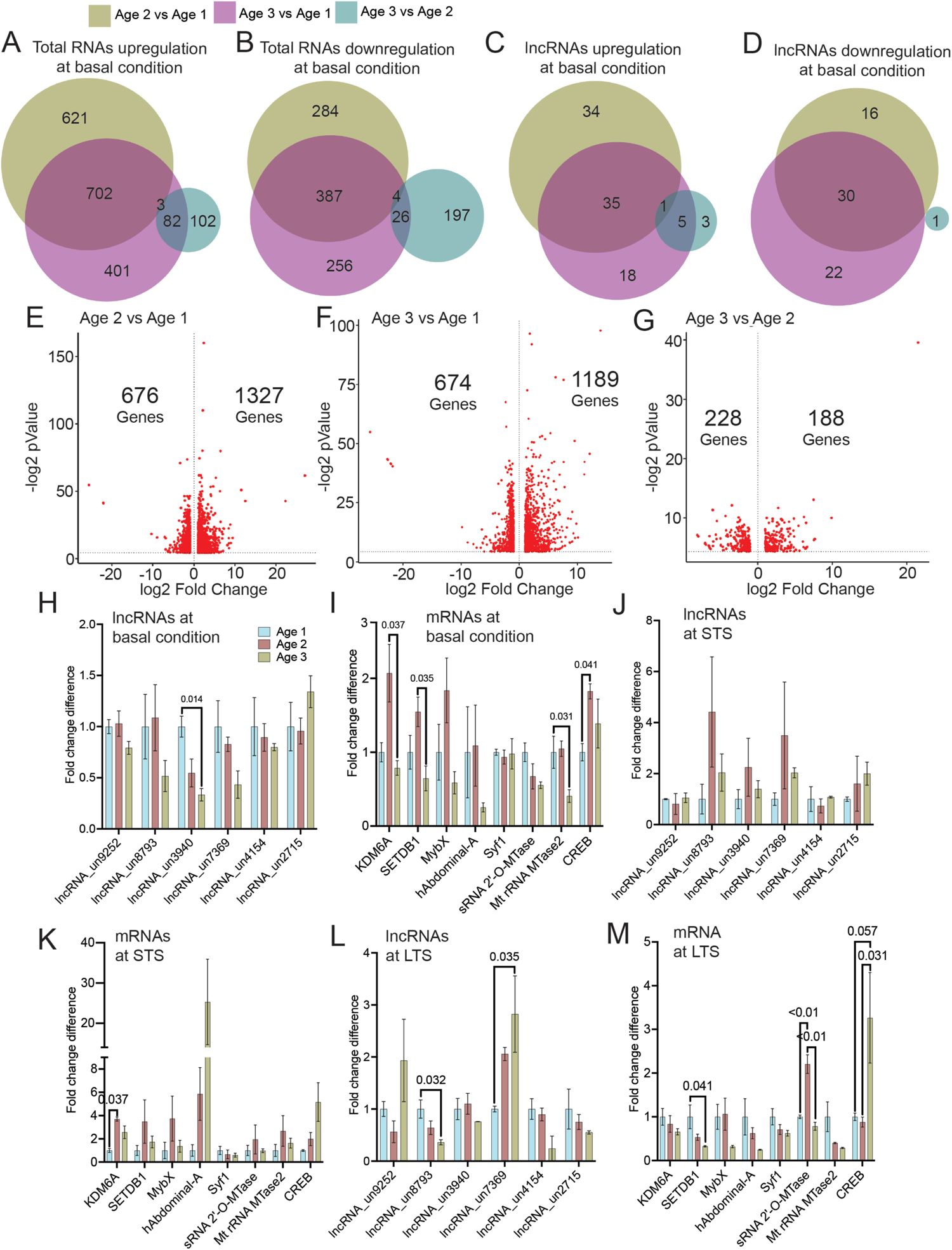
Analysis of aging associated changes in L7MN. RNAseq data from untrained animals (used to generate Figures 2 and 3) were independently compared across the three age groups. Venn diagrams showing comparison of upregulated DEGS (**A),** Down regulated DEGs (**B),** Upregulated lncRNAs (**C),** and down regulated lncRNAs **(D)** (p<0.05)**. E–G.** Differentially expressed genes compared to different age groups are ranked in the volcano plots according to their statistical -log2 p-value (y-axis) and their relative abundance ratio (log2 fold change) between up- and downregulated (x-axis). Red dots indicate significantly regulated genes (false discovery rate, <0.01; s0 = 1; *p*Value <0.05). Re-analysis of qPCR candidates from different age groups (see Figures 2 and 3), **H–I.** at basal condition, **J–K.** in response to short-term sensitization, **L–M.** in response to long-term sensitization. Relative gene expression levels are shown as the mean fold change, with error bars showing the SEM. One-way ANOVA followed by Tukey’s *post hoc* test. N=4, p values are shown in the bar graphs.

To further assess basal level of gene expression changes as well as how aging and learning impacts expression of these candidates across the three age groups, we next re-analyzed qPCR data (Fig. H–M, Supplementary Table S1D). At basal condition, lncRNA_un3940 expression level ∼1.3-fold (p<0.05) significantly reduced at Age 3; on the other hand, at Age 2 KDM6A, SETDB1 levels were found increased compared to Age 3 group. rRNA methyltransferase 2 level in Age 3 group was found significantly lower than Age 1 and 2. CREB level in Age 2 noted higher than Age 1 and Age 3 group. In STS neurons, no significant changes were observed among the age groups, only KDM6A level was significantly higher (∼3.7-fold; *p*<0.05) in Age 2 compared to Age 1 group. Analyzing LTS samples we noted lncRNA_un8793 expression is significantly lower (∼2.8-fold; *p*<0.05) in Age group 2 than Age 1 group. By contrast, lncRNA_7369 level in Age 3 is significantly (∼2.8-fold; *p*<0.05) higher than other age groups. Among LTS samples, mRNAs, SETDB1 showed notable decrease (∼3.1-fold, *p*<0.05) at Age 3 group. sRNA 2’-O-MTase level in Age 2 showed highest (∼2.2-fold, *p*<0.01). CREB level in Age 3 group is ∼3.2-fold higher (*p*= 0.05) compared to Age 1 and 2 group. Take together these results further confirm that aging impairs transcriptional plasticity in L7MN, a key neuron involved in siphon withdrawal reflex.

### STS and LTS regulated genes in L11 and R2 neurons

Our RNAseq analyzes and independent validations in L7MN reveal transcriptome wide changes associated with decline in LTS associated with aging. Further, these analyzes have established that STS and LTS differentially alter the composition of transcriptome in L7MN. We next sought to determine how aging, STS and LTS impact transcriptome of two other identified neurons in the abdominal ganglia, L11MN and a giant cholinergic neuron R2 (Figure 5), that are previously not known to be involved in mediating STS or LTS of siphon withdrawal reflex. R2 is a silent neuron and is part of the *Aplysia* reflex system. During sensitization, *Aplysia* may undergo escape locomotion and the R2 neuron assists in this behavior (*48*). Moreover, gene expression in R2 has been shown to be altered with aging (*48*). L11 is a genital, bursting neuron with projections to the gill and has roles in foot contraction and locomotion in *Aplysia* (*50*). Therefore, we isolated L11MN and R2 from trained and untrained animals and assessed the expression of selected lncRNAs and mRNAs (see Figure 2) in these neurons by qPCR. Importantly, similar to L7MN there is only one L11MN and R2 neuron in the entire animal and are localized in the abdominal ganglia.

**Figure 5.**
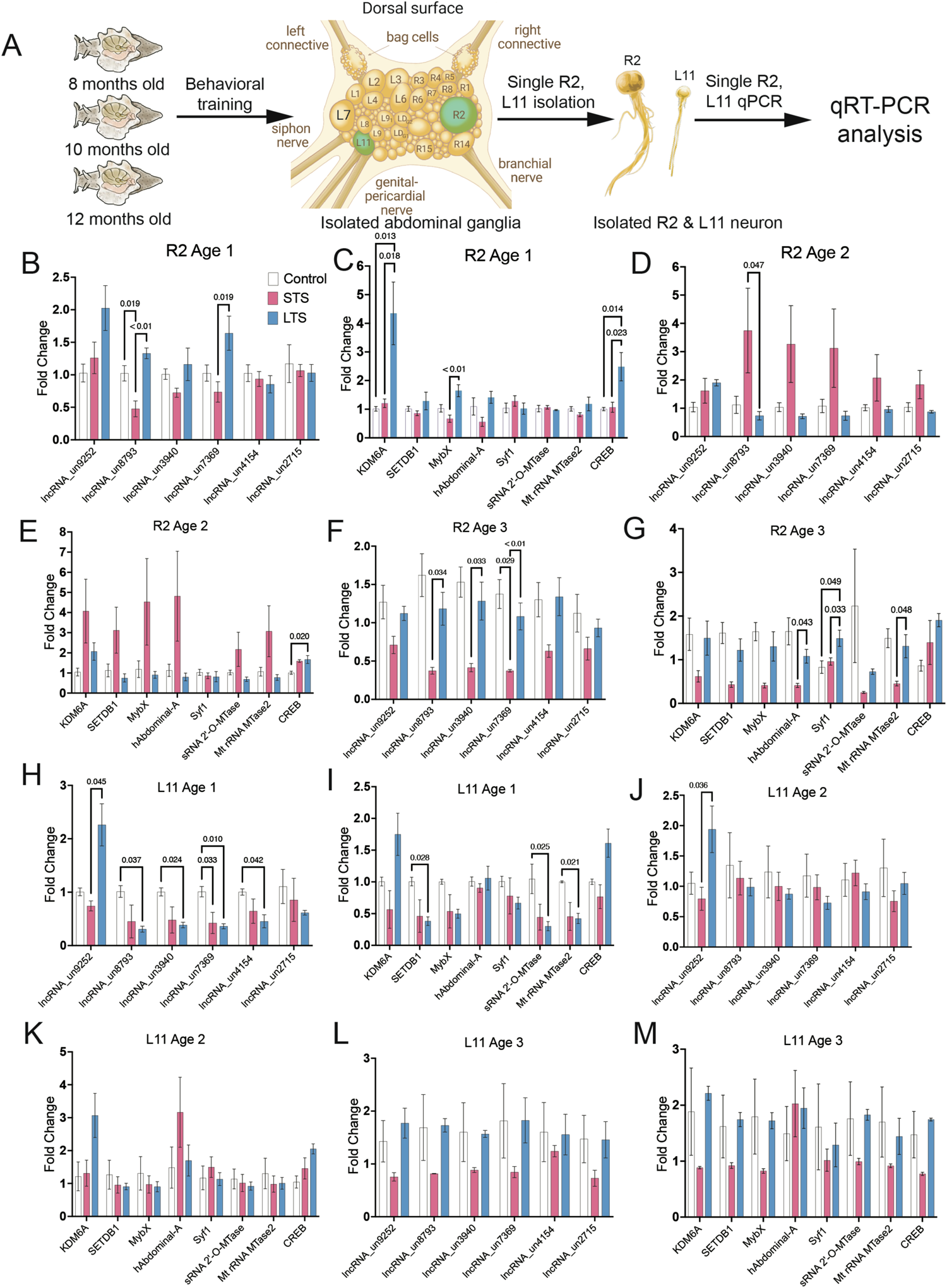
Gene expression analysis of R2 and L11 MN neurons following STS and LTS training. **A.** Schematic representation of the workflow for single R2 and L11 neuron isolation and qPCR analysis from trained (STS and LTS), and untrained control Aplysia from the three age groups. **B–G.** Analysis of qPCR candidates in R2 across different age groups. **H–M.** Analysis of qPCR candidates in L11 across different age groups. Relative gene expression levels are shown as the mean fold change, with error bars showing the SEM. One-way ANOVA followed by Tukey’s *post hoc* test. N= 5, p values are shown in the bar graphs.

Based on the lack of known involvement of L11MN and R2 in sensitization and aging, we anticipated no change in gene expression following training. However, our single L11MN and R2 qPCR show that in response to STS and LTS expression of these genes are altered (up or down regulated) in unique ways (Figure 5) (Supplementary table S1D). LncRNAs 8793 was altered in both STS (down regulated) and LTS (upregulated), 7369 (upregulated in LTS), and mRNAs KGM6A, MyoX and CREB was altered in R2 in age group 1 which became impaired in age group 1 whereas in age group 2 only fewer candidates were altered (lncRNA 8793 and mRNA CREB), but in age group 3, more lncRNAs (8793, 3940, 7369) and mRNAs (hAbdominal A, Syt1, MtrRNA MTase 1) were altered in response to LTS when compared to STS training. By contrast, L11 exhibited recruitment of multiple lncRNAs (five out of 6 lncRNAs studies) and mRNAs (3 out of 8, KDM6A and CREB did not reach significance) in age group 1. These results are similar to L7MN. Interestingly these changes did not persist (except for lncRNA 9252 in age group 2) in age groups 2 and 3. Taken together these results show that lncRNAs and mRNAs are differently modulated in neuron specific manner during aging and learning. Furthermore, these results suggest recruitment of R2 and L11 during STS and LTS learning in addition to L7MN.

### Comparative analysis of expression changes in L7, L11 and R2 neurons suggest neuron specific regulation of learning relevant genes

The finding that some of the genes modulated in L7MN by STS and LTS are also modulated in R2 and L11 led us to examine whether these genes are modulated to the similar extent across these neurons. Significant differences in the fold up or regulation of these candidate genes across these genes would suggest neuron specific modulatory mechanisms. We therefore compared the magnitude of fold changes in these neurons by re-analyzing qPCR data. Data was normalized to corresponding 18s rRNA levels in each neurons (L7MN, L11MN and R2). The analysis across these three neurons identified multiple lncRNAs and mRNAs that are significantly enriched in these neurons in basal levels as well as in response to STS/LTS training. For example, lncRNA 3940, a downregulated in L7MN in response to STS and LTS was found to be enriched L7 MN when compared to R2 and L11 in basal conditions. LncRNA 9252 upregulated in L7MN in response to LTS is also upregulated in L11MN, however, is significantly enriched in L7MN compared to L11MN. Interestingly expression of hAbdominal-A mRNA was enriched in R2 when compared to L7MN and L11MN, STEDB1 was enriched in L7MN compared to L11MN and R2, CREB was enriched in L7MN and L11MN compared to R2 in age group 1 (Supplementary Figures 6 and 7) (Supplementary table S1E).

### Identification of lncRNA-mRNA associations modulated by LTS

We next focused on bioinformatics analysis of lncRNAs to identify lncRNA-mRNA associations modulated by STS and LTS. We first asked whether DEG lncRNAs are enriched in the nucleus or cytoplasm. Nuclear enriched lncRNAs are known to interact transcriptional and epigenetics machinery to modulate gene expression. lncRNAs could potentially modulate expression of RNAs 200 Kb within their locus. Therefore, we first assessed the expression of lncRNAs in nucleus versus cytoplasm fractions and then identified potential regulated RNAs within 200 kb of their loci.

We isolated total RNAs from the abdominal ganglia and checked the expression levels of the top 25 lncRNAs identified from L7MN RNAseq (Supplementary Table S1F) in nuclear and cytoplasmic RNA fractions, and looked for the lncRNAs localized in the nucleus, as they could be potential regulators of transcription (*51*). Systematic screening of these uncharacterized lncRNAs revealed 10 lncRNAs are located in the nucleus (Table 8). Further analysis with the RNAseq dataset, we identified that uncharacterized protein LOC101855924 (LOC101855924) in the vicinity of lncRNA_un8793 (∼4.2 fold upregulated; Age 1 group LTS) and is inversely regulated showed ∼1.2-fold downregulation (Age1 LTS vs STS), TBC1 domain family member 13 (LOC101853398) in the vicinity of lncRNA_un0492 (∼3.2 fold upregulated, Age 2 group LTS) is ∼0.66 fold upregulated (Age2 LTS vs STS). Mucin-5AC (LOC10185663) and golgin subfamily A member 3 (LOC101846517) in the vicinity of lncRNA_un7178 (∼2.57 fold downregulated, Age 2 LTS) and showed ∼1.3-fold and 0.63-fold downregulation (Age2 LTS vs CTRL and Age2 LTS vs STS). Uncharacterized protein LOC101856926 (LOC101856926) and histone H2A-like (LOC101855053) are in the vicinity of lncRNA_un3167 (∼2.02 fold downregulated, Age 2 LTS downregulated) and found ∼1.4 and ∼1.5 fold downregulated (Age2 LTS vs CTRL and Age2 LTS vs STS) (Supplementary Table S1D). Importantly regulation of these lncRNA-mRNA pairs were not observed in Age group 3. Thus, these observations suggest that specific lncRNA-mRNA pairs are recruited during learning but are impaired during aging in L7MN.

## DISCUSSION

Several studies have suggested transcriptome wide changes associated with aging (*12–16*). However little is known about whether aging alters the expression of plasticity relevant genes in neurons mediating LTM. Therefore, identifying learning induced changes in specific components of the transcriptome is a critical step in understanding the impact of aging on LTM (*26, 52, 53*). Identification of transcriptional signatures correlated with aging and learning may pave way for the development of therapeutics targeted to subpopulation of neurons most susceptible to impairments during aging. In this work, we focused on a key neuron (L7MN) involved in two temporally distinct non-associative learning paradigms, short-term sensitization (STS) and long-term sensitization (LTS), to assess the impact of aging on transcriptional plasticity.

Our approach successfully identified STS and LTS induced changes in the long-noncoding and coding transcriptome of L7MN. While all of the DEG lncRNAs identified are uncharacterized, DEG mRNAs are involved in mediating transcriptional, translational, cytoskeletal and synaptic functions. These results are consistent with previous findings on aging associated neuronal changes (*12–16*). Genes differentially modulated by STS and LTS include genes involved in epigenetic and transcriptional regulation (KDM6A and CREB are examples) suggesting that unlike STS, LTS produces long lasting changes in the transcriptional landscape of L7MN. Analysis of STS and LTS modulated genes across the three age groups suggested progressive impairments in their expression with age. In Age group 2 and 3, both STS and LTS induced changes in KDM6A were absent whereas in Age group 2, CREB regulation was absent. In contrast to regulation of CREB in age group 1, there was no difference between in STS and LTS in modulating CREB in age group 3, supporting the lack of sensitization in age group 3.

Previous studies have documented lncRNA expression changes associated with aging (*54–56*). However, modulation of learning relevant lncRNAs by aging remains poorly understood. We observed notable changes in the expression of specific lncRNAs and their predicted targets in response to LTS. Our analysis shows that LTS modulated lncRNAs also exhibited impairments during aging. Importantly, unlike other age groups modulation of predicted lncRNA-mRNA pairs were absent in age group 3. Taken together, these results establish progressive impairments in transcriptional plasticity during aging.

We wondered about the specificity of STS/LTS induced changes in the transcriptional landscape of L7MN. Specifically, we wanted to know whether STS/LTS induced genes are also modulated in two other neurons-L11MN and R2 that do not have a known function in STS or LTS of siphon withdrawal. Our qPCR analysis of candidate genes (lncRNAs and mRNAs identified in L7MN) suggested transcriptional modulation of specific genes by STS/LTS in L11 and R2. Compared to R2, these changes in L11MN were more pronounced.

It’s been shown that L11MN undergoes regulation by both paracrine and autocrine diffusible factors from sensory neurons following synapse formation (*57*). Moreover, L11MN has roles in Aplysia foot contraction and locomotion (*50*). While it is not known whether L11MN participate in LTS, the discovery that L11MN could be modulated by diffusible factors suggests its potential role in LTS. Potentially, activation of sensory neurons and interneurons during LTS training produces diffusible factor/s that modify signaling in L11MN. However, this needs to be experimentally tested. The transcriptional changes observed in L11MN and R2 in response to STS/LTS training suggest that these neurons might have a role in mediating LTS. Together, these analysis suggest neuron specific changes during LTS (lncRNA 9252 and 7369 in both L7MN and L11, KDM6A and CREB in L7MN and R2). In all three of these neurons, in age group 3 transcriptional modulation of specific genes is either completely lacking or altered (including both upregulation and down regulation of specific transcripts) suggesting severe impairments in transcriptional plasticity in senescent animals.

Importantly, all three of these neurons are located in the same ganglion, the abdominal ganglion of *Aplysia* CNS, yet they exhibit different trajectories of aging. Collectively, our data indicate that aging induced changes might not be identical in all cell types, which is in line with previous findings (*5, 6, 8, 22*). Together, these observations underscore the significance of single neuron and neural circuit based assessments of aging to identify specific deficits produced by aging. We identified both qualitative and quantitative changes in the coding and noncoding transcriptomes during aging. Further we found that aging occurs progressively in a cell specific manner and that these changes likely underlie aging associated cognitive decline. We were able to validate a subset of transcriptional changes. However, a large number of lncRNAs and mRNAs remains to be characterized. While our work has established transcriptomic correlates of aging associated impairments in LTS, future work will be required to assess the functions and mechanisms of differentially regulated lncRNAs and mRNAs, as well as the role of L11MN and R2 in LTS.

We anticipate that data presented in this manuscript will serve as a resource for the neuroscience community and for those who study the biology of aging and learning. For example, our lncRNA data could facilitate future studies aimed at determining the role of the non-coding transcriptome in modulating plasticity and aging. Integrating these studies with functional assays may reveal how noncoding and coding transcriptomes interact for neuronal plasticity and how aging impacts their interaction and function. Understanding how transcriptomic changes in individual neurons modulate specific learning and LTM will be vital for obtaining novel mechanistic insights underlying aging associated cognitive decline and for developing therapeutics.

## MATERIALS AND METHODS

### Animals

#### Aging cohorts

Two cohorts of animals were reared and maintained at the National Resource of Aplysia at the University of Miami’s Rosenstiel School of Marine, Atmospheric, and Earth Science. Cohort one (group 1) hatched on February 2, 2019. Cohort two (group 2) hatched on March 1, 2019. Animals were reared at 15C and fed red algae ad libitum before training began. Behavioral training, including sensitization with 1 or 4 tail shocks (*33, 58*) and no shock controls was performed on animals at 8 (Age 1), 10 (Age 2), and 12 months (Age 3) of life.

#### Before the pre-test

Thirty animals from each cohort were selected for each age group to investigate age-related memory deficits in *Aplysia*. If possible, active animals with similar body sizes were chosen for training. Because the siphon withdrawal reflex was used to measure long-term memory capacity, animals with larger siphons were preferred. Animals were selected for training based on appearance, weighed, and placed in individual cages for one week before the pre-test. Algae access was restricted three days before the pre-test. Observations such as egg mass formation, abnormal locomotion, animal physical appearance, and weight were noted throughout the experiment. *Aplysia* body mass increases during development but declines after sexual maturity and aging.

#### Behavior training Short-term and long-term sensitization

##### Pre-test

On day one of training, a paintbrush bristle was used as a non-noxious stimulus (touch) to elicit siphon withdrawal. The duration or latency of withdrawal (SWRL) from the time of the stimulus to the beginning of relaxation was recorded by a blind observer. Each animal received four touches. The animal’s average SWRL or pre-test value was calculated and used to group the animals for training so that each group’s average SWRL pre-test values was similar.

##### Behavioral training

Five groups were used to investigate age-related learning deficits: two groups for behavioral measurements (B) following four shocks for long-term sensitization (LTS) or no shock control, and three groups for single-cell isolation (SCI) and RNA analysis following either one shock for short-term sensitization (STS), four shocks for LTS, or no shock control. Day two of training included mock tail shocks for the control groups and either a single tail shock or four tail shocks separated by 30 minutes for the sensitization groups (*34, 59*). For the single-cell isolation groups, animals’ abdominal ganglia were dissected one hour after training, and single neurons (L7, L11, and R2) were isolated for RNA analyses.

##### Test

On day three, the behavioral groups’ long-term memory was tested. Four siphon touches were elicited, and the average SWRL was compared to the average pre-test value as a measure of training retention.

### Isolation of L7MN, L11 MN and R2 for RNA extraction

To investigate the transcriptional dynamics at a single neuron level, we isolated the L7MN, L11MN and R2 motor neurons from the abdominal ganglia from the STS and LTS-trained sea slugs. Following 1h after the last shock, the abdominal ganglia were dissected from the animals and single neurons were collected as described by Akhmedov et al. (2013 (*60*)). The total RNA was extracted using the Arcturus™ PicoPure™ RNA Isolation Kit (Applied Biosystems), and subjected to total and small RNA sequencing. See supplementary Table S1A for details of the batch, and neurons used for RNA isolation and analysis.

One hour after behavior training, Aplysia were injected with isotonic MgCl_2_ for 5-10min (equivalent to 30-35% of the animal’s body weight). Following the methodology protocol from Akhmedov et al. (2013), the abdominal ganglia (with long, intact L. and R. connective nerves, and as long as possible siphon, genital-pericardial and branchial nerves) were isolated and treated with 0.1-0.3% protease in artificial seawater (ASW) for ∼1-2 hours, depending on body weight. After digestion, ganglia were pinned in a Sylgard Silicone chamber and perfused with ASW, desheathed, and the target neurons were identified as described in Akhmedov et al. (2013). Areas around the L7, L11, and R2 neurons were cleared, and neighboring neurons were removed to ensure the isolation of single cells. Then, 100% ethanol was perfused over the ganglia to petrify the neurons, and target neurons were individually isolated with forceps and placed along the wall of a frozen non-stick 1.7mL microfuge tube on dry ice.

The axon length and thickness of R2, L7, and L11 differ. R2 axons project from the left abdominal ganglion to the ipsilateral, contralateral pedal, and pleural ganglia (Moroz & Kohn, 2010). The L7 axons project to the siphon, genital-pericardial, and branchial nerves (*61*). L11 has many branches from its axons. While each target neuron’s average thickness and length are unique, we established a grading system (A-E) to classify the length of the axons from each isolated neuron. Neurons with relatively exceptionally long axons were an “A.” Neurons with relatively long axons were classified as “B.” Neurons with short axons were graded as a “C.” Neurons with a small segment of axon were considered “D.” Lastly, neurons with only the cell body isolated were considered “E.” Only L7 “A” and “B” isolated neurons were considered for RNA sequencing.

### RNAseq Analysis

RNAs were isolated from single L7MN using the Arctus LCM RNA isolation kit, and the quality of RNAs was assessed using a bioanalyzer. We obtained 20 ng/ul RNAs (total 10 ul eluted RNA from one microdissected cell body) that we used for RNAseq (Clontech SMART-Seq Ultra Low Input RNA kit) in Scripps Florida Genomics Core (Currently known as The Herbert Wertheim UF Scripps Institute for Biomedical Innovation & Technology). After removal of ribosomal RNAs using a custom kit developed at the Scripps Genomics Core, RNAs were sequenced using Hiseq500. In this experiment, we obtained ∼ 20 million reads per sample (n=4-6 for each condition). After quality control, the reads were mapped to the Aplysia genome (seahare-NCBI-aplcal3.0).

RNAseq analysis are carried out by Maryland Genomics, Institute for Genome Sciences, UMSOM. Illumina libraries are mapped to the *Aplysia californica* reference, NCBI RefSeq accession GCF_000002075.1, using HiSat2 v2.0.4, using default mismatch parameters. Read counts for each annotated gene are calculated using HTSeq. The DESeq2 Bioconductor package (v1.5.24) is used to estimate dispersion, to normalize read counts by library size, to generate the counts per million for each gene, and to determine differentially expressed genes between experiment and control samples. Differentially expressed transcripts with a raw p-value ≤ 0.05 and a minimum 1.5X fold-change between groups were used for downstream analyses. RNAseq was deposited to NCBI and can be accessed (GEO accession number: GSE234983)

### Quantitative real-time PCR

Following our previously stated protocols (*58, 62, 63*) quantitative real-time PCR (qRT-PCR) analyses were conducted to validate the RNAseq data. After 1h of behavioral training, L7, L11 and R2 neurons were collected from the abdominal ganglia as described in the previous section from all age groups. Using the Arcturus™ PicoPure™ RNA Isolation Kit, total RNA was extracted from the single neurons individually, and cDNA was prepared using qScript cDNA supermix. Aplysia 18S rRNA reference gene is used to normalization. Relative quantification of each transcript was done following the 2^ΔΔCt^ method (*64*).

### lncRNA target analysis

To analyze the cytoplasmic or nuclear localization of the lncRNAs detected (Supplementary table S1F) in the RNAseq experiment, cytoplasmic or nuclear fractionation of RNA was isolated from the abdominal ganglia using Norgen Biotek Corp Cytoplasmic & Nuclear RNA Purification Kit following manufacturers protocol. cDNAs were generated from the purified RNA using qScript cDNA supermix were used in the qPCR analysis. We next focused on nuclear enriched lncRNAs and manually searched for potential RNAs transcribed 200 kb upstream or downstream of the loci of candidate lncRNAs (potential cis-regulated RNAs) by manually searching *Aplysia* genome sequences. We then selected predicted cis-targets and examined whether they are among the DEGs identified from the RNAseq data from L7MN. lncRNAs identified from RNAseq and also within 200 kb of lncRNA loci were considered as potential targets of candidate lncRNAs.

### Statistical analyses

Statistical analyses were conducted in R and Prism 9. All data used for preparing Figures and corresponding statistical analyses are available in the Supplementary Table file. Behavior data was analyzed by using a 3 way ANOVA. Unpaired two-tailed Student’s t test and one-way ANOVA followed by Dunnett’s post hoc test were used unless indicated otherwise. The results are graphically represented as the mean ± standard error of the mean (SEM) throughout the text, unless otherwise stated. N represents the number of independent samples for each experiment.

## AUTHOR CONTRIBUTIONS

SVP designed the project with inputs from RDH; PG, DS help set up aging cohorts and maintained them; LF, and MCS provided all infrastructural resources and guidance regarding setting up aging cohorts; KB optimized single neuron isolations, carried out all behavior analyses, prepared and interpreted behavior data, and isolated L7MN, L11MN and R2 neurons; AS carried out all L7MN qPCR analyses and prepared all Figures; BLR and SLV analyzed R2 and L11 neurons by qPCRs; CM conducted all bioinformatics analyses with help from AS and AM; SVP, AS, BLR interpreted results; SVP and AS wrote the paper, and SP and RDH revised the manuscript based on inputs from authors.

## Supporting information

https://www.dropbox.com/scl/fi/u48axdc5ggu0rc2higlnq/Badal-et-al-Supplementary-Figures-2023.pdf?dl=0&rlkey=96sebpqzqridyhoh1gzbkwxtu

https://www.dropbox.com/scl/fi/nwunzng8dlv6jh4bi1tgb/Supplementary-Tables-Badal-et-al-Aplysia-aging-2023.xlsx?dl=0&rlkey=r0fgx1717lxm26u1gt5gkm10f

## ACKNOWLEDGEMENTS

We gratefully acknowledge the funding support from NSF (Grant 1453799) to SVP, and NIH (1R21AG055049, 1R01MH119541 and 1R01MH118444) to SVP, 1F31MH127958-01A1 to KKB, and P40OD010952 to MCS. We thank Dr. Pabalu Karunadharma, Director of Scripps Florida Genomics Core, for helping us with developing ribodepletion tools for *Aplysia* and optimization of single neuron RNAseq; Vani Addepalli and Marina Perez, Undergraduate students at Florida Atlantic University, for helping with setting up initial behavior and neuron isolation experiments.

## CONFLICT OF INTEREST STATEMENT

The authors declare no conflicts of interests.

## DATA AVAILABILITY STATEMENT

All the RNAseq data are available from NCBI, accession number: GSE234983.

## SUPPORTING INFORMATION

Supplementary Figures: 7

Supplementary Tables: 6

**Supplementary Figure 1. Analyses of nuclear versus cytoplasmic localization of the lncRNAs. A.** qPCR analyses of relative expression levels of 25 lncRNAs distribution. with error bars showing the SEM. Statistical analyses were conducted by paired two-tailed student’s T-test. **B.** Bioinformatic analyses based on NCBI and RNAseq data to identify lncRNA and *cis-*regulated transcripts. Relative gene expression levels shown as the mean fold change, p values are shown in bar graphs.

**Supplementary Figure 2. Heat maps showing the normalized and scaled expression values of the top 50 differentially expressed genes when ranked by p-value.** The color gradient from green to red represents high to low expression levels across the samples. The genes are ordered by hierarchical clustering using Euclidean distance and complete clustering method while the samples are ordered by condition. **A–C.** Age 2 STS versus Control, Age 2 LTS versus Control, Age 2 LTS versus STS respectively. **D–F.** Age 3 STS versus Control, Age 3 LTS versus Control, Age 3 LTS versus STS respectively.

**Supplementary Figure 3. qPCR analyses of RNAseq candidates in R2 across different age groups.** This is a re-analysis of data from Figure 5. qPCR analyses of relative expression levels of lncRNAs and mRNAs in R2 : **A–B.** Basal condition, **C–D.** short-term sensitization, **E–F.** long-term sensitization. Relative gene expression levels are exhibited as the mean fold change, with error bars showing the SEM. One-way ANOVA followed by Tukey’s *post hoc* test. N=5, p-Values are indicated in bar graphs.

**Supplementary Figure 4. qPCR analyses of RNAseq candidates in L11 across different age groups.** This is a re-analysis of data from Figure 5. qPCR analyses of relative expression levels of lncRNAs and mRNAs in L11: **A–B.** Basal condition, **C–D.** short-term sensitization, **E–F.** long-term sensitization. Relative gene expression levels are exhibited as the mean fold change, with error bars showing the SEM. One-way ANOVA followed by Tukey’s *post hoc* test. N=5, p-Values are indicated in bar graphs.

**Supplementary Figure 5. Analysis of fold change of gene expression across L7, R2 and L11 in Age group 1**. This is a re-analysis of data from Figures 1 and 5. **A–B.** Basal condition, **C–D.** short-term sensitization, **E–F.** long-term sensitization. Relative gene expression levels are exhibited as the mean fold change, with error bars showing the SEM. One-way ANOVA followed by Tukey’s *post hoc* test. N=5, p-Values are indicated in bar graphs.

**Supplementary Figure 6. Analysis of fold change of gene expression across L7, R2 and L11 in Age group 2**. This is a re-analysis of data from Figures 3-5. **A–B.** Basal condition, **C–D.** short-term sensitization, **E–F.** long-term sensitization. Relative gene expression levels are exhibited as the mean fold change, with error bars showing the SEM. One-way ANOVA followed by Tukey’s *post hoc* test. N=5, p-Values are indicated in bar graphs.

**Supplementary Figure 7. Analysis of fold change of gene expression across L7, R2 and L11 in Age group 3**. This is a re-analysis of data from Figures 3-5. **A–B.** Basal condition, **C–D.** short-term sensitization, **E–F.** long-term sensitization. Relative gene expression levels are exhibited as the mean fold change, with error bars showing the SEM. One-way ANOVA followed by Tukey’s *post hoc* test. N=5, p-Values are indicated in bar graphs.

## Notes

### Competing Interest Statement

The authors have declared no competing interest.

