## Supplementary figures and images for "Single neuron analysis of aging associated changes in learning reveals progressive impairments in transcriptional plasticity"

### https://www.dropbox.com/scl/fi/u48axdc5ggu0rc2higlnq/Badal-et-al-Supplementary-Figures-2023.pdf?dl=0&rlkey=96sebpqzqridyhoh1gzbkwxtu

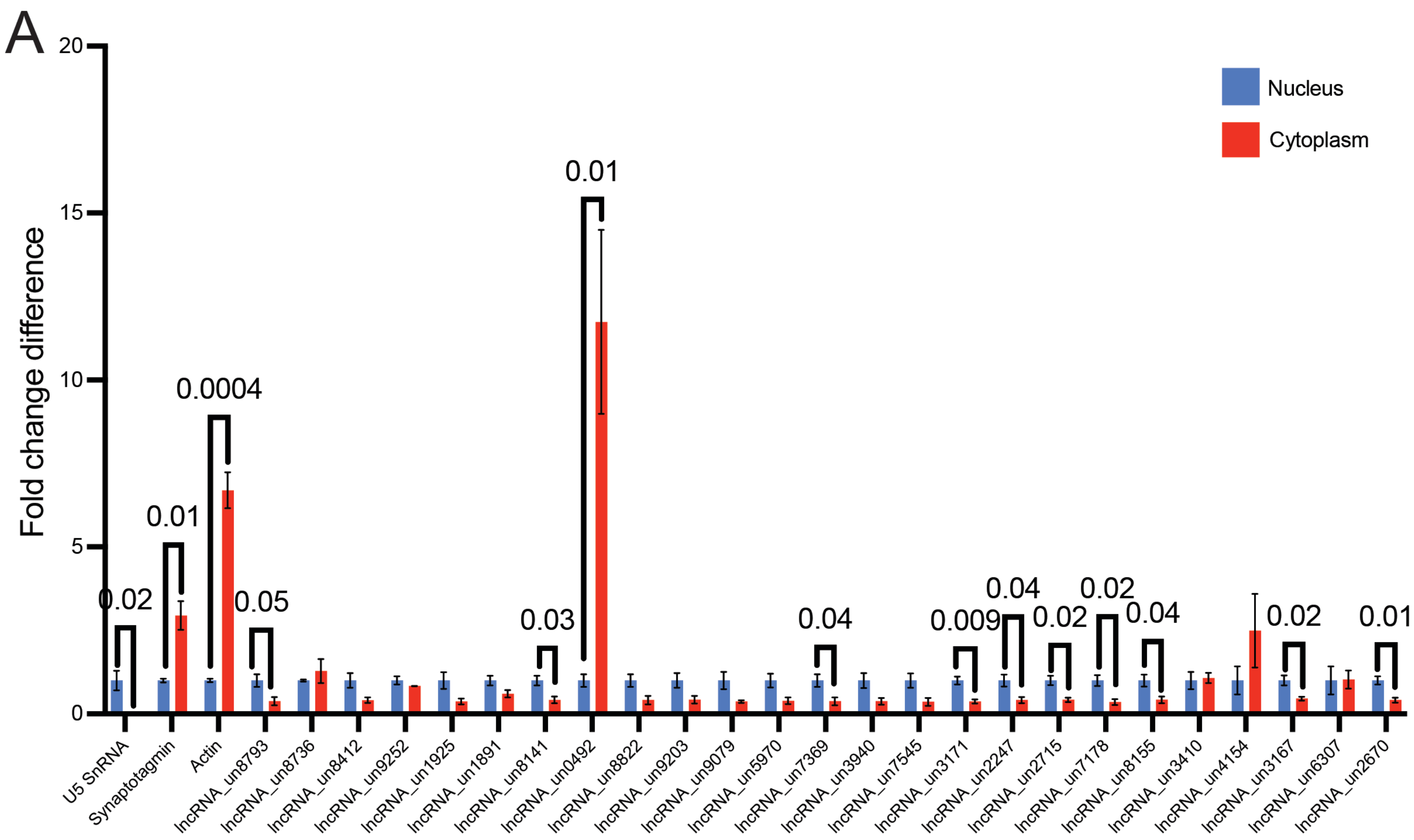

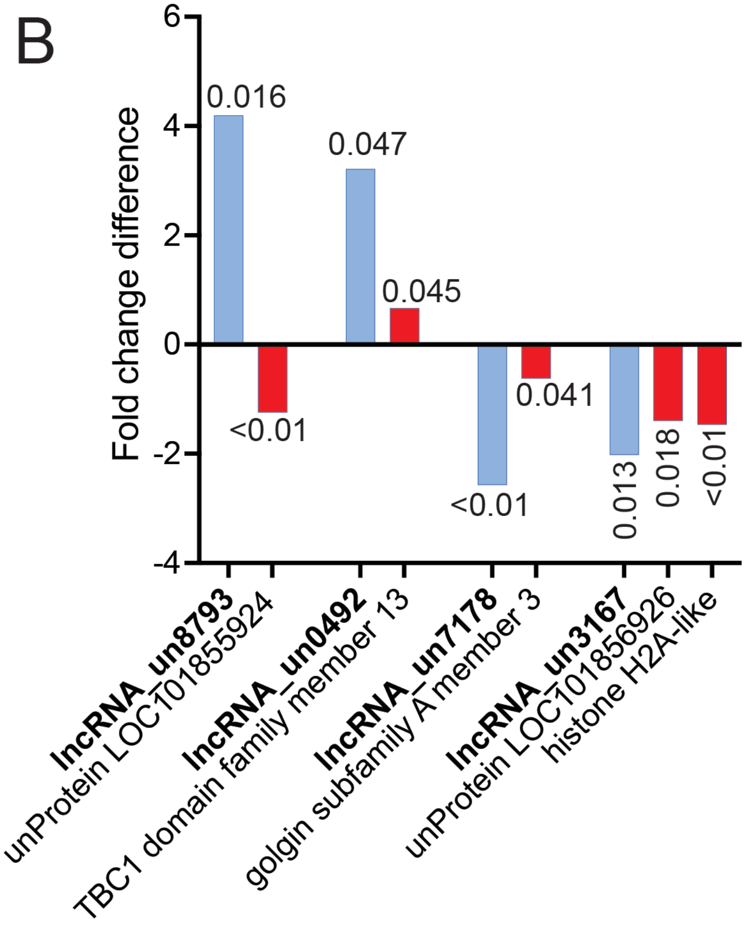

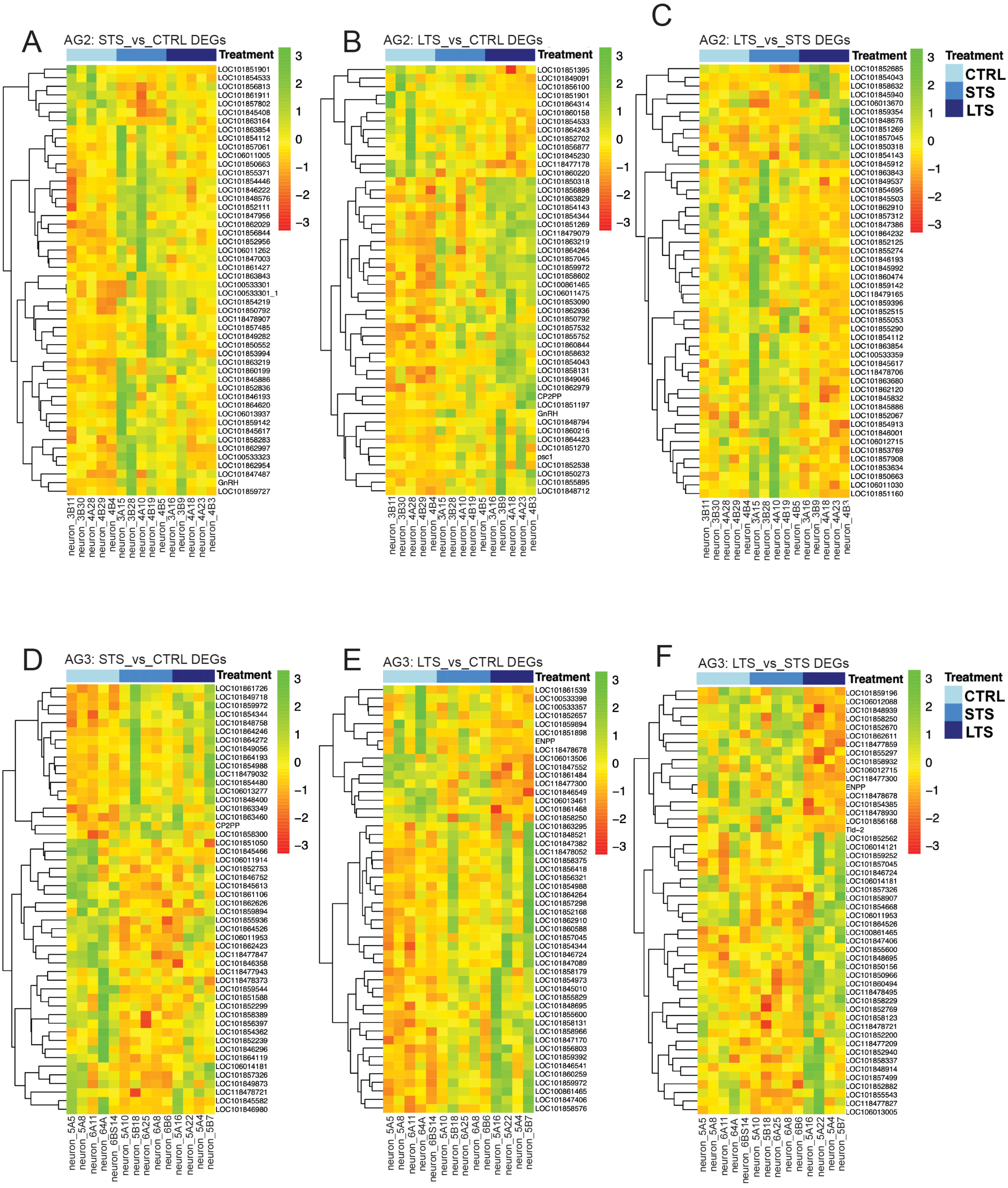

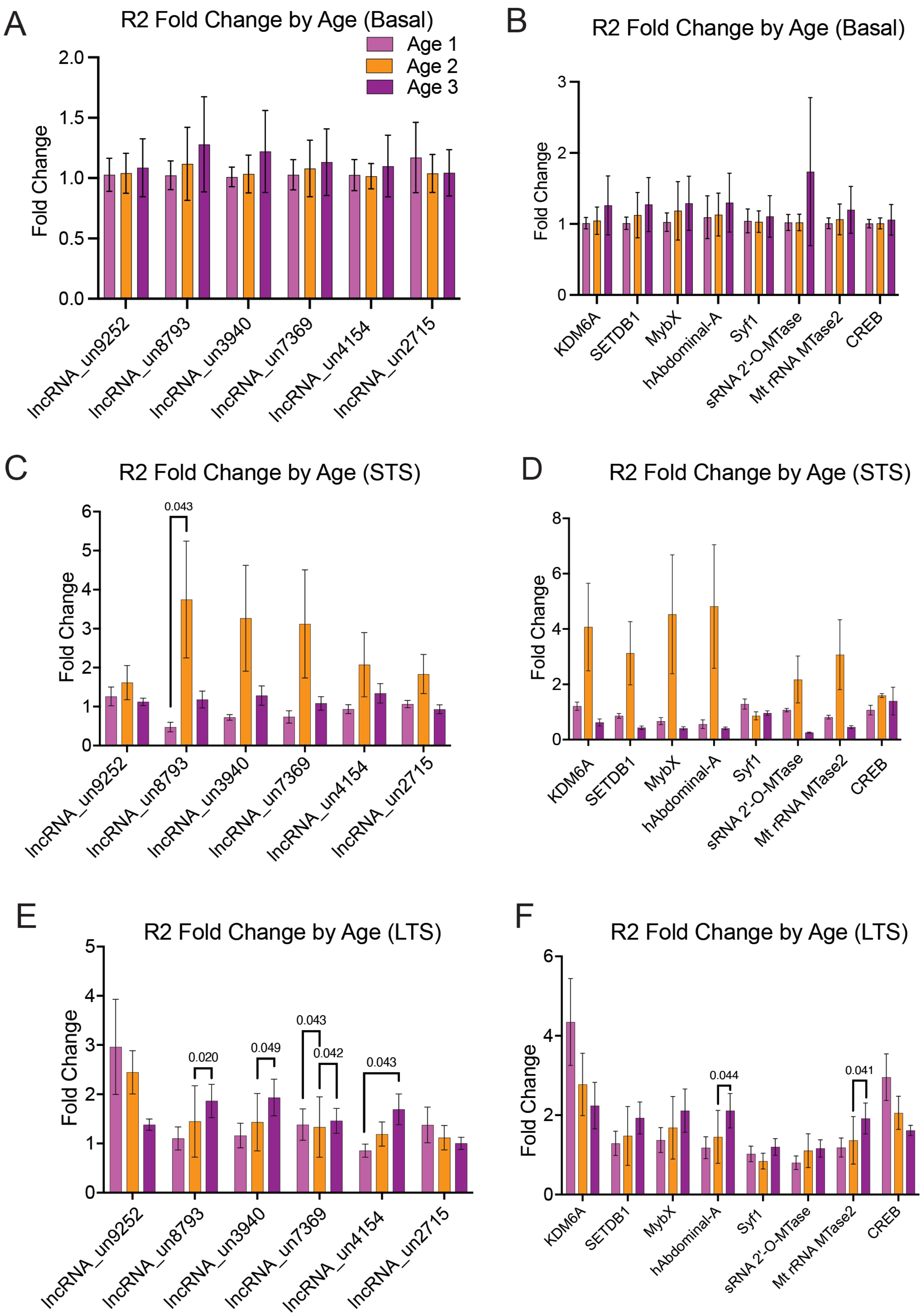

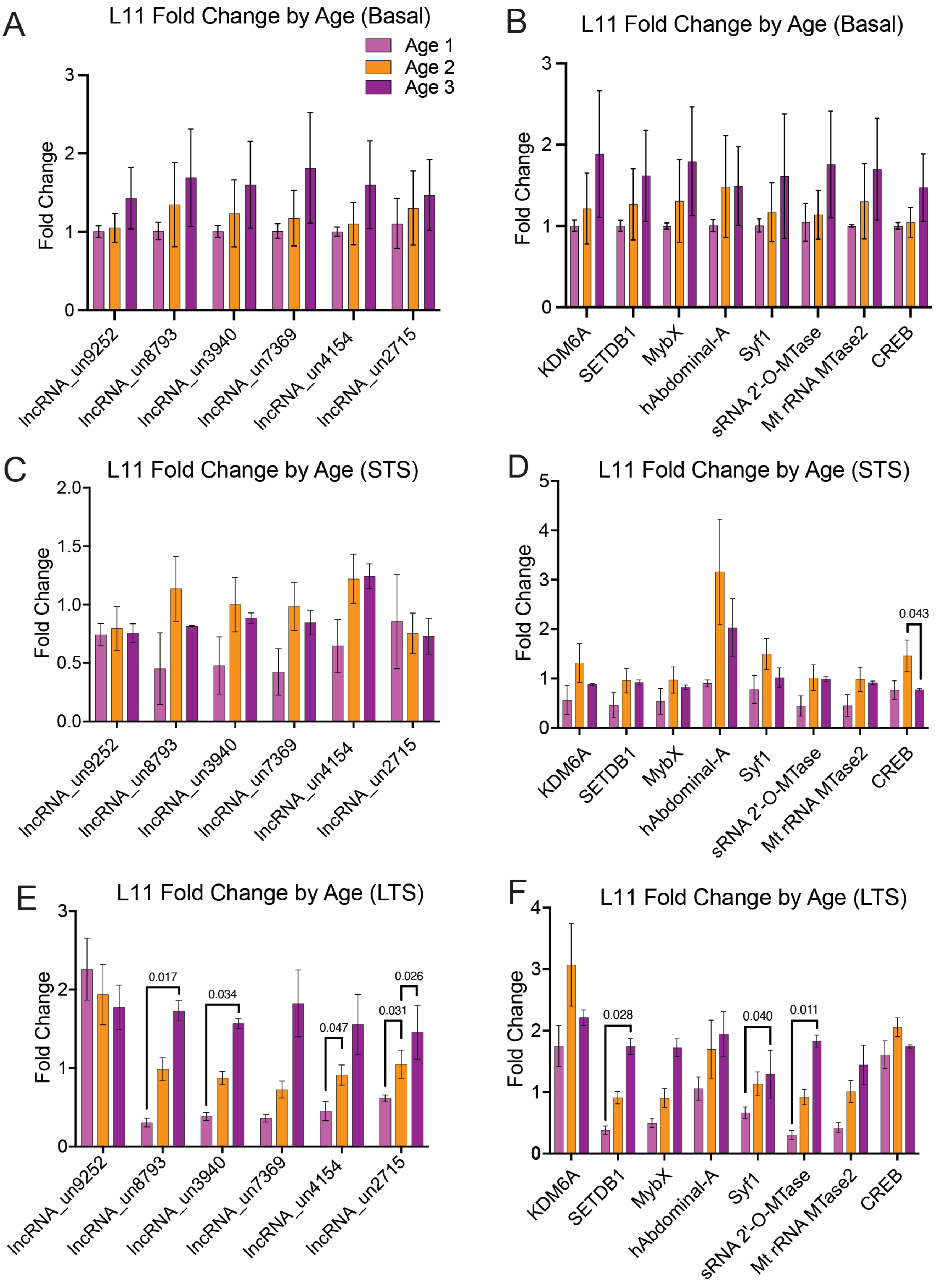

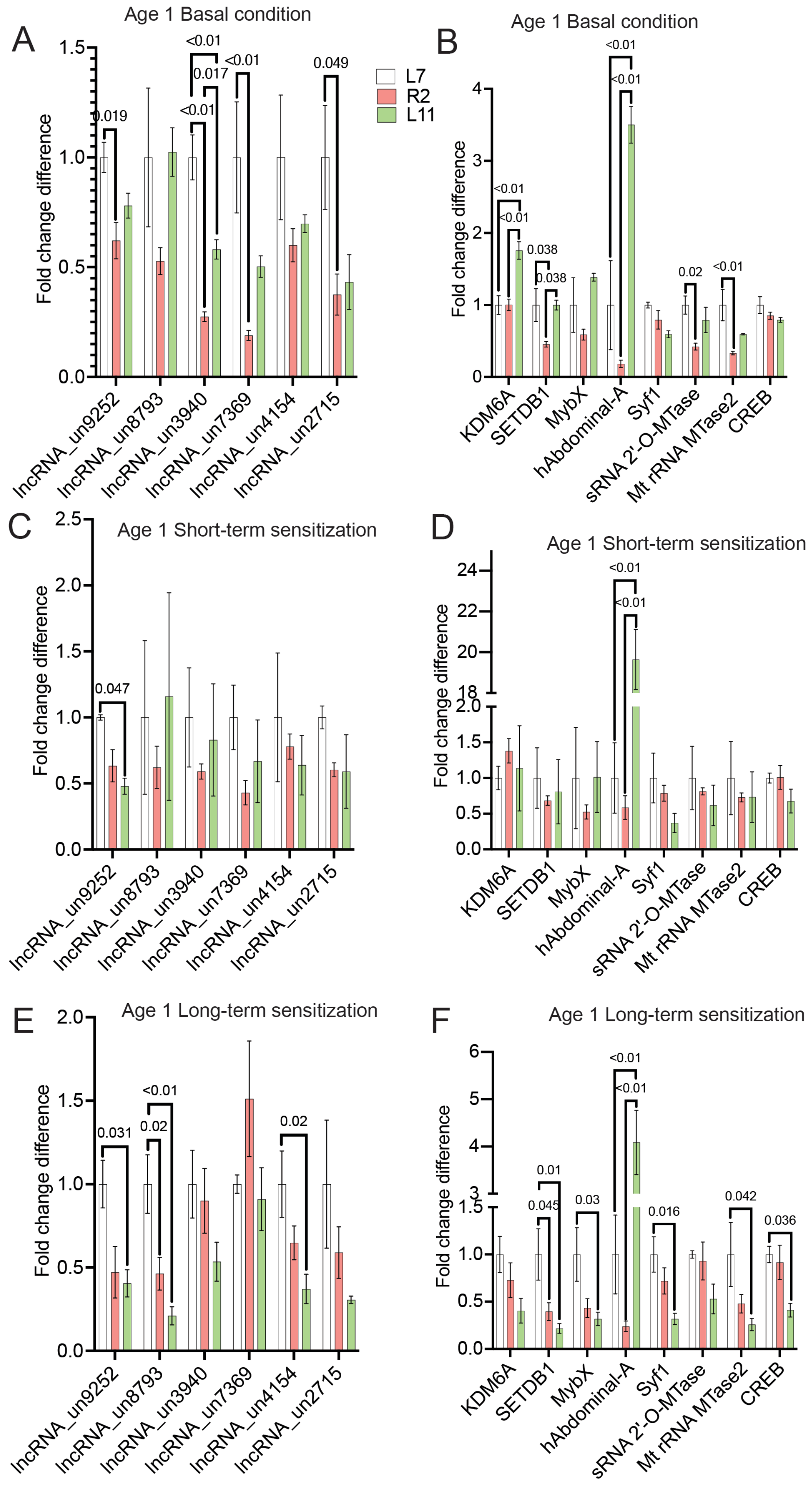

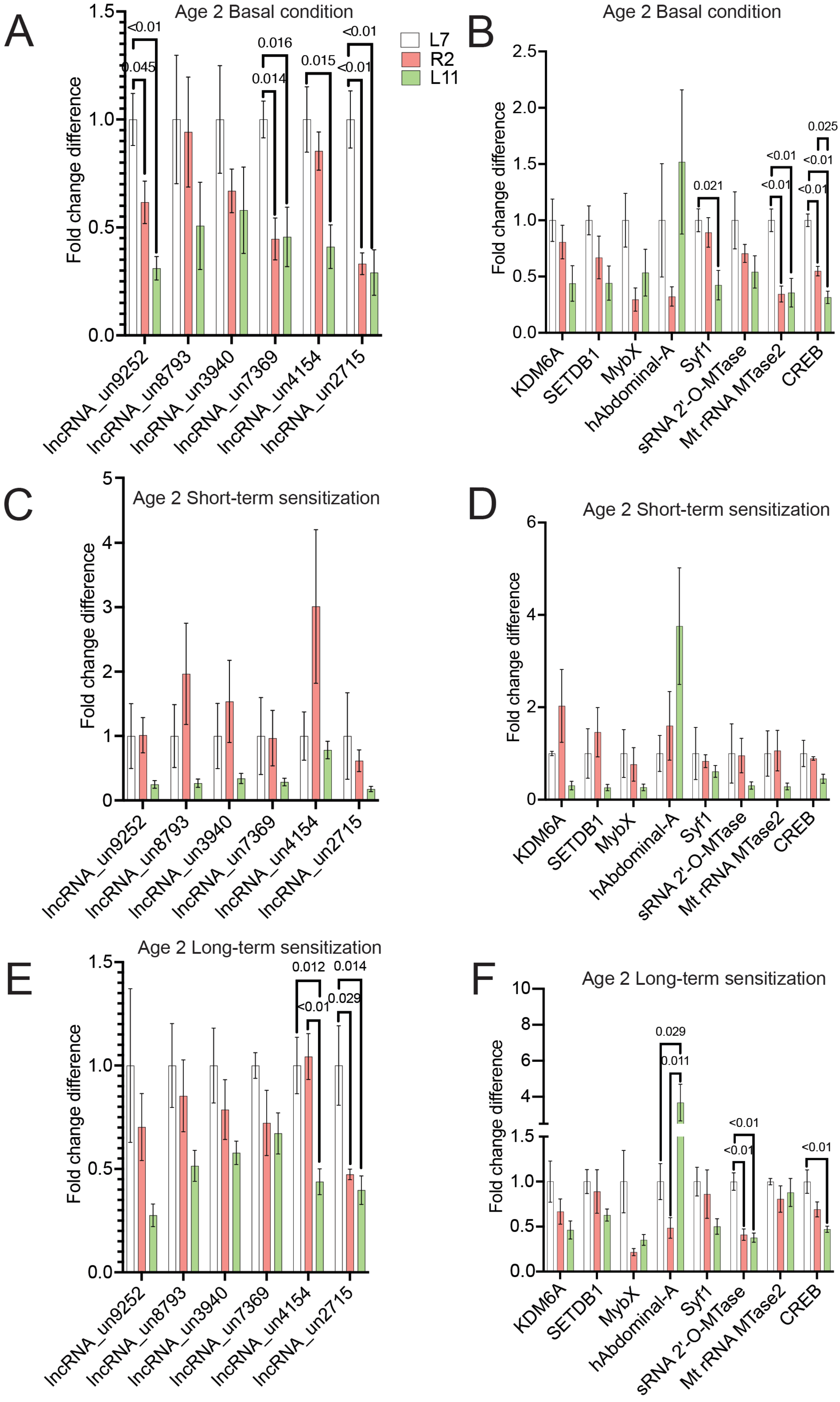

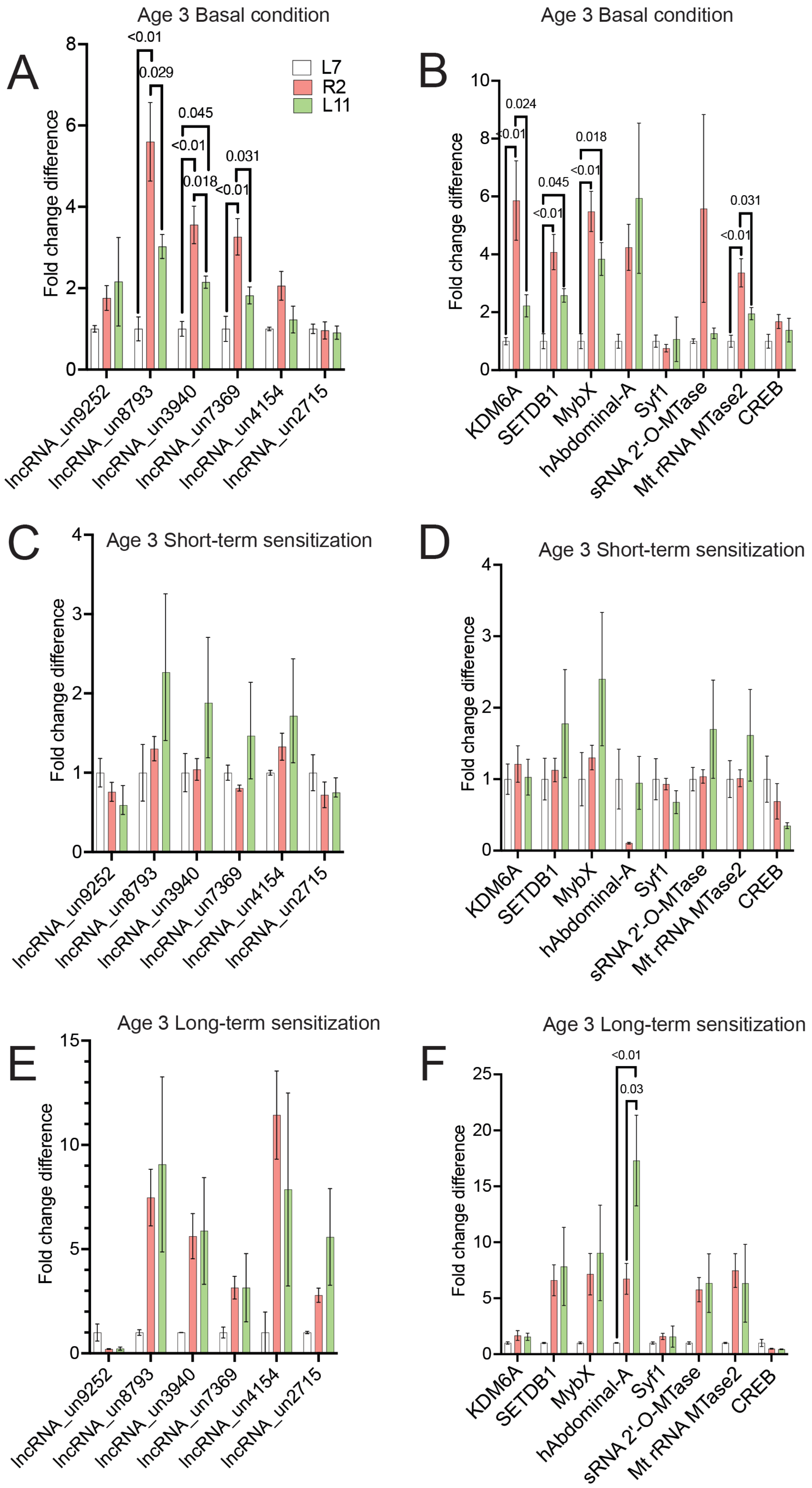
